# SARS-CoV-2 evolution and patient immunological history shape the breadth and potency of antibody-mediated immunity

**DOI:** 10.1101/2022.05.06.490867

**Authors:** Maria Manali, Laura A. Bissett, Julien A. R. Amat, Nicola Logan, Sam Scott, Ellen C. Hughes, William T. Harvey, Richard Orton, Emma C. Thomson, Rory N. Gunson, Mafalda Viana, Brian Willett, Pablo R Murcia

**Affiliations:** MRC-University of Glasgow Centre for Virus Research. Glasgow, United Kingdom; School of Veterinary Medicine, College of Medical, Veterinary and Life Sciences, University of Glasgow, Glasgow, United Kingdom; West of Scotland Specialist Virology Centre, NHS Greater Glasgow and Clyde, Glasgow, United Kingdom; Institute of Biodiversity, Animal Health and Comparative Medicine, University of Glasgow, Glasgow, United Kingdom

**Keywords:** SARS-CoV-2, immunological history, antibody-mediated immunity, virus neutralisation, virus evolution

## Abstract

Since the emergence of SARS-CoV-2, humans have been exposed to distinct SARS-CoV-2 antigens, either by infection with different variants, and/or vaccination. Population immunity is thus highly heterogeneous, but the impact of such heterogeneity on the effectiveness and breadth of the antibody-mediated response is unclear. We measured antibody-mediated neutralisation responses against SARS-CoV-2_Wuhan_, SARS-CoV-2α, SARS-CoV-2δ and SARS-CoV-2ο pseudoviruses using sera from patients with distinct immunological histories, including naive, vaccinated, infected with SARS-CoV-2_Wuhan_, SARS-CoV-2α or SARS-CoV-2δ, and vaccinated/infected individuals. We show that the breadth and potency of the antibody-mediated response is influenced by the number, the variant, and the nature (infection or vaccination) of exposures, and that individuals with mixed immunity acquired by vaccination and natural exposure exhibit the broadest and most potent responses. Our results suggest that the interplay between host immunity and SARS-CoV-2 evolution will shape the antigenicity and subsequent transmission dynamics of SARS-CoV-2, with important implications for future vaccine design.

**Author Summary:** Neutralising antibodies provide protection against viruses and are generated because of vaccination or prior infections. The main target of anti-SARS-CoV-2 neutralising antibodies is a protein called Spike, which decorates the viral particle and mediates viral entry into cells. As SARS-CoV-2 evolves, mutations accumulate in the spike protein, allowing the virus to escape antibody-mediated immunity and decreasing vaccine effectiveness. Multiple SARS-CoV-2 variants have appeared since the start of the COVID-19 pandemic, causing various waves of infection through the population and infecting-in some cases-people that had been previously infected or vaccinated. Since the antibody response is highly specific, individuals infected with different variants are likely to have different repertoires of neutralising antibodies. We studied the breadth and potency of the antibody-mediated response against different SARS-CoV-2 variants using sera from vaccinated people as well as from people infected with different variants. We show that potency of the antibody response against different SARS-CoV-2 variants depends on the particular variant that infected each person, the exposure type (infection or vaccination) and the number and order of exposures. Our study provides insight into the interplay between virus evolution and immunity, as well as important information for the development of better vaccination strategies.

## Introduction

Severe acute respiratory syndrome coronavirus 2 (SARS-CoV-2) emerged in December 2019[1] causing the largest pandemic of the XXI century. Since the start of the pandemic, different viral lineages emerged, exhibiting highly dynamic transmission patterns[2–5]. This is illustrated by the epidemiology of COVID-19 in the United Kingdom since the first introduction of SARS-CoV-2 in late February 2020, when the Wuhan strain (SARS-CoV-2_W_) was first reported. A D614G variant of the Wuhan strain circulated almost exclusively until September of the same year, when the alpha strain (SARS-CoV-2α) appeared, initially in the Southeast of England[2]. By December 2020, SARS-CoV-2α was the dominating variant. At this point in time, the UK started a COVID-19 vaccination program that took place with unprecedent pace: by July 1^st^,2021, ~47 million people (mainly adults) had received at least one vaccine dose[6]. However, during this period, a new variant (delta, SARS-CoV-2δ) emerged in India, reaching the UK in April 2021, and quickly became the most prevalent lineage, until November 2021, when the omicron variant (SARS-CoV-2ο) was introduced and quickly replaced SARS-CoV-2δ. Over a period of approximately 2 years (February 2020 to March 2022) the UK population experienced four COVID-19 pandemic waves, each of them caused by a different SARS-CoV-2 variant (Wuhan, alpha, delta and omicron). In addition, SARS-CoV-2δ and SARS-CoV-2ο infections have been reported in individuals that had been previously vaccinated or infected by preceding variants[7, 8]. As a result, population immunity against SARS-CoV-2 is likely to be highly heterogeneous. The impact of such immunological heterogeneity on SARS-CoV-2 fitness is far from clear. As antibody-mediated immunity is considered a correlate of protection[9, 10], identifying the factors that affect the humoral immune response is key for COVID-19 preparedness and to design more effective vaccines. Our overall objective was to quantify the breadth and potency of antibodies elicited by different immune histories against distinct SARS-CoV-2 variants. To this end, we measured antibody-mediated immunity against SARS-CoV-2_W_, SARS-CoV-2α, SARS-CoV-2δ, and SARS-CoV-2ο using convalescent serum samples from the Glasgow patient population collected between March 31^st^, 2020, and September 22^nd^, 2021. Our sampling strategy captured the complex immunological landscape described above and included sera from naive, vaccinated, infected, as well as vaccinated and infected individuals. Importantly, by combining patient metadata (date of positive PCR) with virus genomic epidemiology (prevalence of circulating lineages over time) we were able to select confidently serum samples from patients exposed to three major variants that circulated in the UK (Wuhan, alpha and delta).

## Results

A schematic description of the study is shown in Fig 1. Serum samples (n=353) from biobanked material that had been collected for serological surveillance studies[11] were selected. The immunological history of each patient at the time of sampling was compiled based on serological status using an ELISA assay that tested for SARS-CoV-2 spike 1 (S1) and receptor binding domain (S-RBD)[11] together with metadata associated with each clinical specimen. Associated metadata consisted of date of serum collection, SARS-CoV-2 PCR status (including date and result of diagnosis) and vaccination status (including date of vaccination and number of doses). Samples were initially classified in four broad groups: *naive* (N, 30 samples), *vaccinated* (V, 55 samples) *infected* (I, 91 samples) and *infected and vaccinated* (I_V_, 177 samples). Further, sera from the I and I_V_ group were stratified based on the infecting variant as *infected-Wuhan* (I_W_, 37 samples), *infected-alpha* (Iα, 39 samples), *infected-delta* (Iδ, 15 samples), *infected-Wuhan-vaccinated* (I_WV_, 60 samples), *infected-alpha-vaccinated* (Iα_V_, 69 samples), and *infected-delta-vaccinated* (Iδ_V_, 48 samples). The date of PCR confirmation and the prevalence of each variant at the time of diagnosis was used to infer the most likely infecting variant (see materials and methods and S1 Fig). All serum samples were processed as follows: first they were tested using a multiplex electrochemiluminescence assay against the spike (S), and nucleocapsid (N) proteins to quantify antibody concentrations. Further, serum samples were subjected to virus neutralisation assays (VNAs)[11] at a fixed dilution (1:50) using HIV (SARS-CoV-2) pseudotypes carrying the S glycoprotein of either SARS-CoV-2_W_, SARS-CoV-2α, SARS-CoV-2δ, or SARS-CoV-2ο. Every sample that displayed 50% neutralisation to at least one SARS-CoV-2 variant was subject to antibody titration as previously described [12].

**Figure 1.**
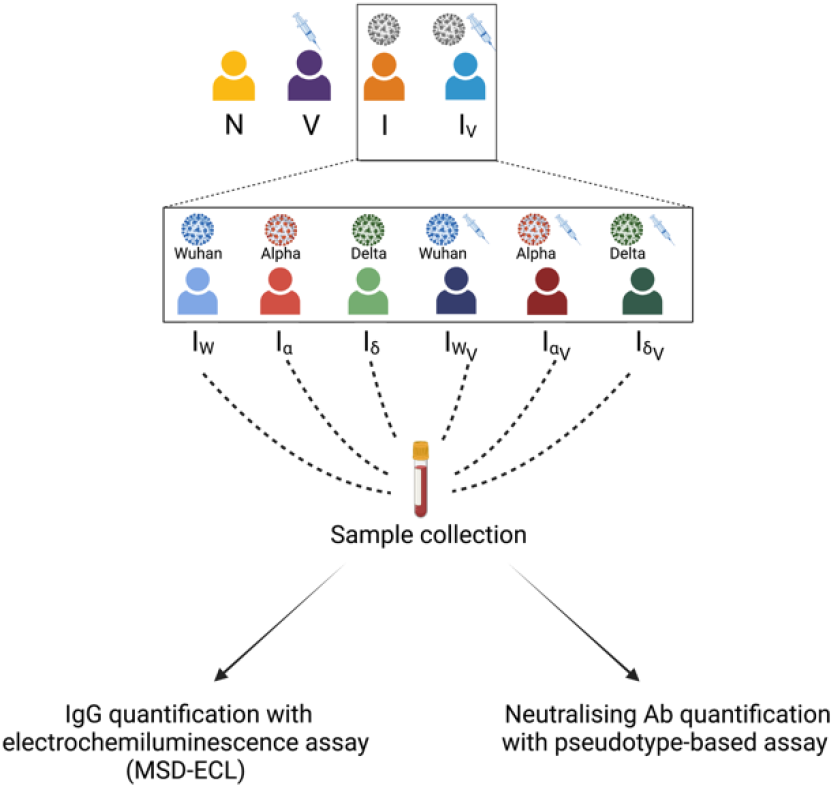
Schematic representation of the study design. Coloured silhouettes represent different patient groups (N= naive, V= vaccinated, I= infected, I_V_= infected and vaccinated). I and I_V_ were further stratified based on the infecting strain (I_W_= infected with SARS-CoV-2_W_, Iα= infected with SARS-CoV-2α, Iδ= infected with SARS-CoV-2δ, I_WV_= vaccinated and infected with SARS-CoV-2_W_, Iα_V_= vaccinated and infected with SARS-CoV-2α, Iδ_V_= vaccinated and infected with SARS-CoV-2δ). Serum samples were tested for the presence of IgG antibodies against SARS-CoV-2 S and N, and also tested in virus neutralisation assays using pseudotyped viruses carrying the S glycoprotein of specific SARS-CoV-2 variants (see methods).

Quantification of S and N antibody levels for each group of patients is shown in Fig 2. As expected, sera from patients in the naive group (neither vaccinated nor infected) exhibited the lowest levels of anti-S antibodies because they had not been exposed to the Spike antigen of SARS-CoV-2. Patients that had been infected displayed higher levels of anti-S antibodies than vaccinated ones, whereas both were significantly lower to those observed in sera from patients that had been infected and vaccinated. Sera from patients that had been infected possessed higher levels of anti-N antibodies than those that had been infected and vaccinated (Fig 2). Of note, vaccinated patients had lower levels of anti-N than naive individuals. Overall, these results are consistent with previous reports suggesting that exposure to SARS-CoV-2 antigens by vaccination and infection results in higher levels of anti-SARS-CoV-2 antibodies than vaccination alone [13–15].

**Figure 2.**
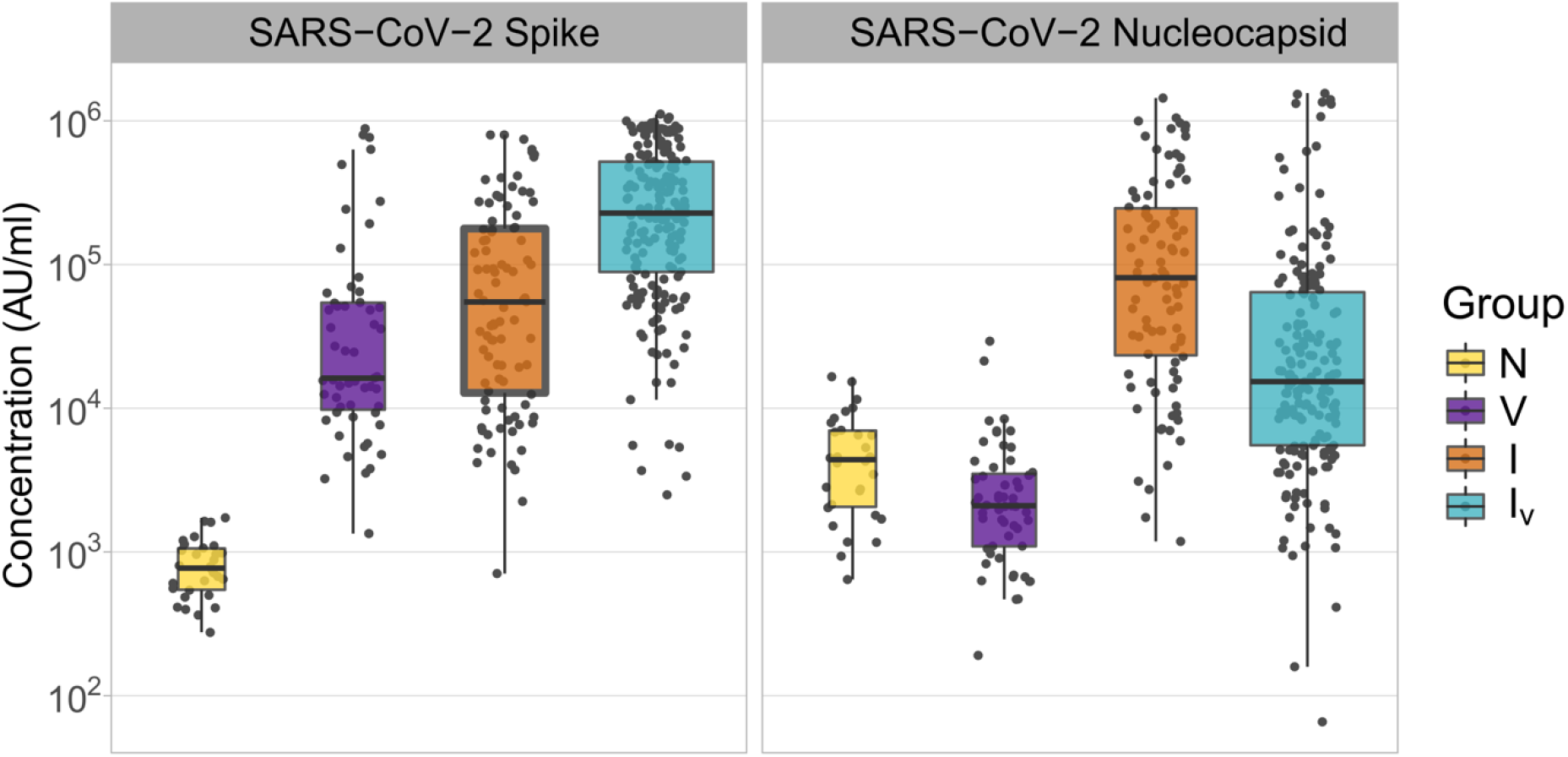
Concentrations of SARS-CoV-2 Spike and Nucleocapsid antibodies in samples derived from patients with different histories of SARS-CoV-2 exposure. The name of the antigens is shown at the top of each panel. Patient groups are defined as N: naïve (yellow); V: vaccinated (purple); I: infected (orange); I_V_: infected and vaccinated (cyan). Antibody concentrations are shown in MSD arbitrary units/ml. Boxplots displayed the interquartile range and median values.

We next measured the neutralisation activity of each serum sample at a fixed dilution against SARS-CoV-2_W_, SARS-CoV-2α, SARS-CoV-2δ, and SARS-CoV-2ο using virus pseudotypes. The efficiency of neutralisation varied depending on the SARS-CoV-2 variant tested (Fig 3A), and the immunological history of the patients (Fig 3B). When the chronological order of appearance of each variant is considered, a pattern of neutralisation reduction consistent with antigenic drift emerges. This is illustrated by the ladder-like distribution of the median percentage neutralisation (Fig 3A) and becomes even more evident when neutralisation levels are compared between SARS-CoV-2ο and all the other variants, as the former, more evolved S, is neutralised less effectively. A similar trend of neutralisation reduction is observed between SARS-CoV-2_W_ and SARS-CoV-2δ in the V and I_V_ groups (Fig 3A). We also observed that virus neutralisation efficiency increases depending on the number and type of exposures, irrespective of the variant tested (Fig 3B). As a result, the I_V_ group exhibited the highest neutralisation values against all variants (Fig 3B).

**Figure 3.**
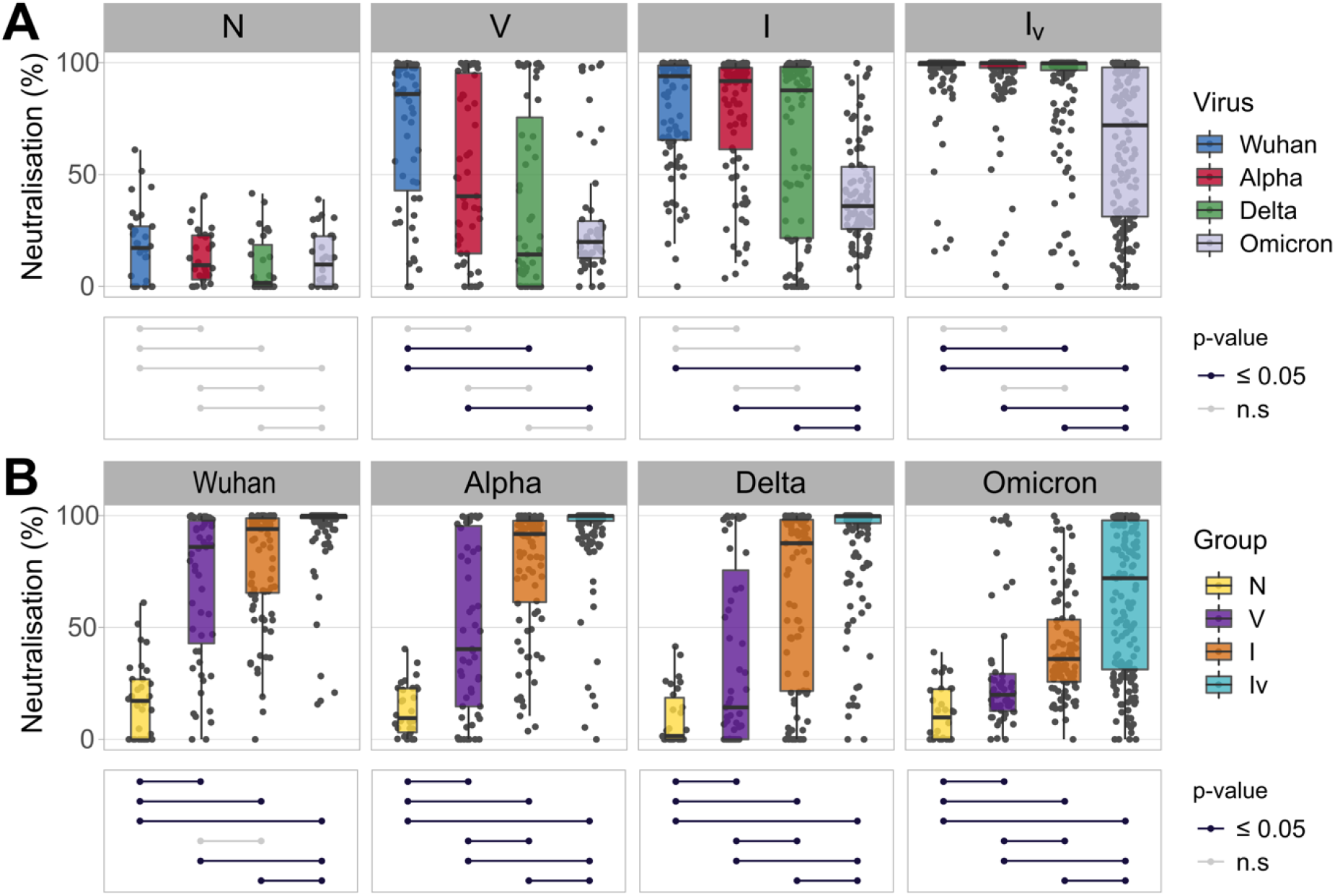
Neutralising responses elicited against pseudotyped viruses carrying the S protein of different SARS-CoV-2 variants according to patient exposure history to SARS-CoV-2. (A) Sera from patients were grouped based on immunological histories (N: naive; V: vaccinated; I: infected; I_V_: infected and vaccinated). Neutralising activity was measured using Wuhan (blue), Alpha (red), Delta (green) and Omicron (grey) spike glycoprotein bearing HIV(SARS-CoV-2) pseudotypes and plotted per patient group (A) and per SARS-CoV-2 S variant (B). Neutralisation was measured at a fixed dilution (1:50). Each point represents the mean of two replicates. Boxplots displayed the interquartile range and median values. Significance levels between patient groups or pseudotyped viruses were tested using pairwise Wilcoxon test, and are shown in bottom panels as connected dotplots.

To quantify more accurately the neutralising potency of the antibody-mediated response among the four broad groups (N, V, I, I_V_), we titrated neutralising antibodies against each variant. Consistent with our previous results, neutralising titres significantly decreased as SARS-CoV-2 evolved (Fig 4A). Indeed, the aforementioned “ladder-like” effect was even more evident. Also consistent with our previous results, the number and type of antigen exposure events had a significant impact on virus neutralisation titres (Fig 4B). Patients derived from the I_V_ group displayed significantly higher neutralising antibody titres compared to every other group across all variants (Fig 4B). In turn, infected patients exhibited variable titres against each variant when compared to vaccinated patients: for example, differences between these two groups were non-significant when SARS-CoV-2_W_, SARS-CoV-2α and SARS-CoV-2δ were compared. However, patients from the V group displayed significantly higher antibody titres against SARS-CoV-2ο, albeit neutralisation efficiency was still very low. Collectively, these results suggest that SARS-CoV-2 antigenic evolution is directional (SARS-CoV-2 evolved to escape antibody-mediated immunity) and that the number and type of exposure events affect the breadth and potency of the antibody-mediated response.

**Figure 4.**
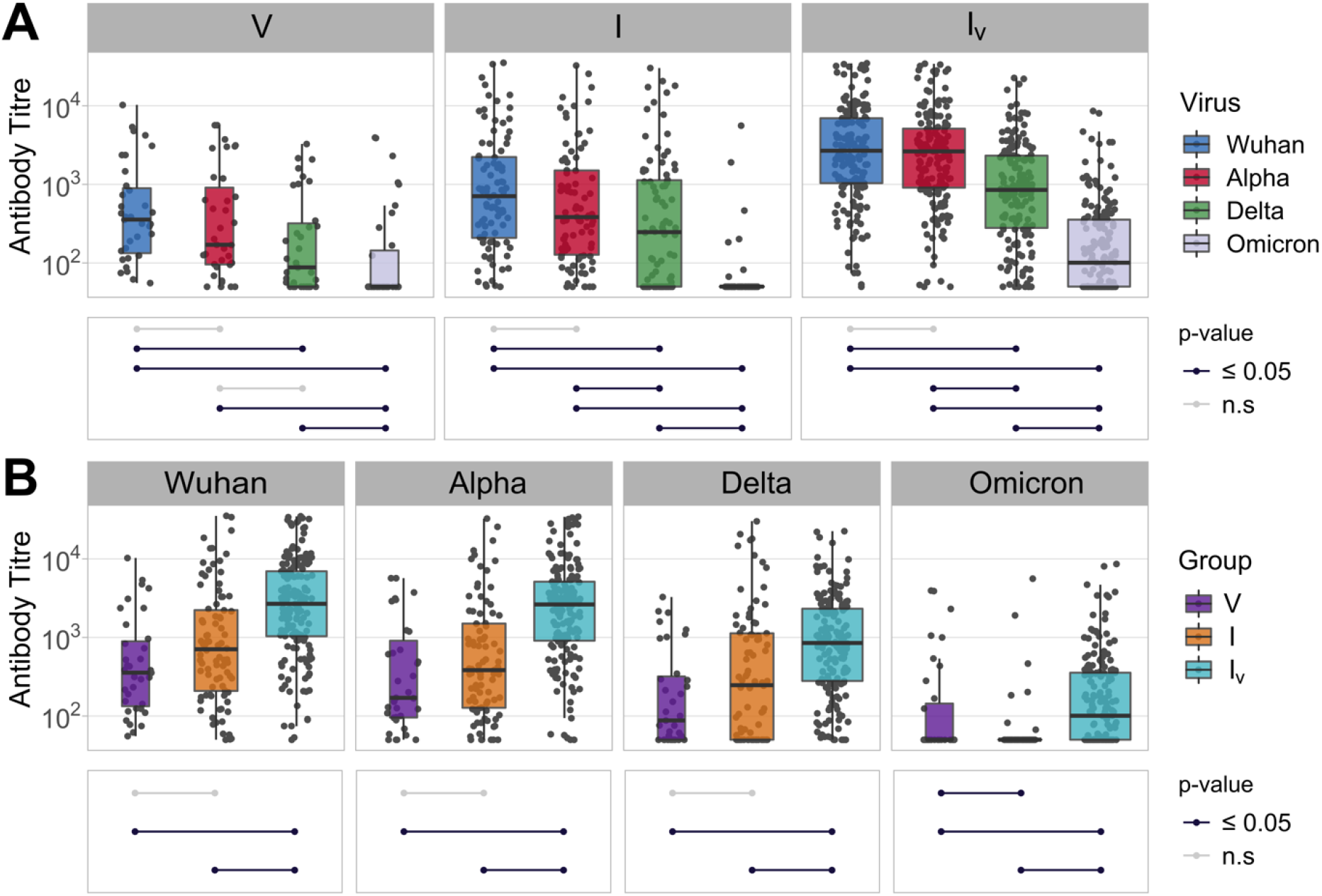
Neutralising antibody titres against SARS-CoV-2 variants in sera from patients with different histories of SARS-CoV-2 exposure (V: vaccinated (purple); I: infected (orange); I_V_: infected and vaccinated (cyan)). Neutralising activity was measured using Wuhan (blue), Alpha (red), Delta (green) and Omicron (grey) spike glycoprotein bearing HIV(SARS-CoV-2) pseudotypes and plotted per patient group (A) and per SARS-CoV-2 S variant (B). Each point represents the mean of three replicates. Boxplots displayed the interquartile range and median values. Significance levels between patient groups or pseudotyped viruses were tested using pairwise Wilcoxon test, and are shown in bottom panels as connected dotplots.

To understand better how humoral immunity is affected by the antigenicity of SARS-CoV-2 variants, we stratified the I and I_V_ groups according to the strains that had infected the patients. This analysis revealed a trend consistent with homologous immunity as sera from patients that had been infected with SARS-CoV-2_W_, SARS-CoV-2α or SARS-CoV-2δ displayed highest potency against their infecting variants, albeit differences were not always statistically significant (Fig 5A). Notably, patients that had been vaccinated and infected with SARS-CoV-2δ showed overall the highest neutralisation potency against all variants (Fig 5B). As this group of patients had been exposed to the most phylogenetically distant antigens (SARS-CoV-2_W_ by vaccination and SARS-CoV-2δ by infection), this result suggests not only that heterologous exposure results in a broader and more effective humoral response but also that the level of antigenic differences between variants affects the potency of the antibody-mediated response. In addition, we observed some differences among patients that had been vaccinated and infected with each SARS-CoV-2 variant: for example, patients that had been infected with SARS-CoV-2α exhibited lower neutralisation efficiency against SARS-CoV-2δ than against SARS-CoV-2_W_ or SARS-CoV-2α (Fig 5A). Titration of neutralising antibodies enabled us to quantify neutralisation biases towards specific SARS-CoV-2 variants. Generally, patients infected by specific variants exhibited significantly different neutralising titres against other SARS-CoV-2 variants (Fig 6A and B) and this effect was also evident among vaccinated and infected patients. For example, patients infected with SARS-CoV-2δ exhibited high levels of neutralising antibodies against SARS-CoV-2δ but significantly lower titres against all other variants (Fig 6A), whereas patients infected with SARS-CoV-2_W_ or SARS-CoV-2α displayed similar levels of neutralising antibodies against SARS-CoV-2_W_ and SARS-CoV-2α but lower levels against SARS-CoV-2δ and even lower against SARS-CoV-2ο (Fig 6A). Overall, titres in patients that had been infected only were lower to those measured in patients that had been infected and vaccinated (Fig 6A and B). The only exception was observed in sera from patients infected with SARS-CoV-2δ, whose neutralising antibody titres against the homologous antigen was similar in vaccinated and infected patients (Fig 6B). Neutralising antibody responses seemed to display immunological preferences: for example, patients that had been vaccinated and infected with SARS-CoV-2δ exhibited significantly higher neutralisation levels against SARS-CoV-2_W_ (the vaccine variant) than SARS-CoV-2δ (the infecting variant), suggesting the stimulation of an anamnestic response. In contrast, patients that had been vaccinated but infected with SARS-CoV-2α showed similar neutralising antibody titres against SARS-CoV-2_W_ and SARS-CoV-2α. Of note, when neutralising antibody titres were compared across all patient groups, those vaccinated and infected with SARS-CoV-2δ or infected with SARS-CoV-2_W_ and then vaccinated, displayed the highest titres against all variants (Fig 6B), consistent with the notion that immunity conferred via infection *and* vaccination results in broader and more potent humoral responses.

**Figure 5.**
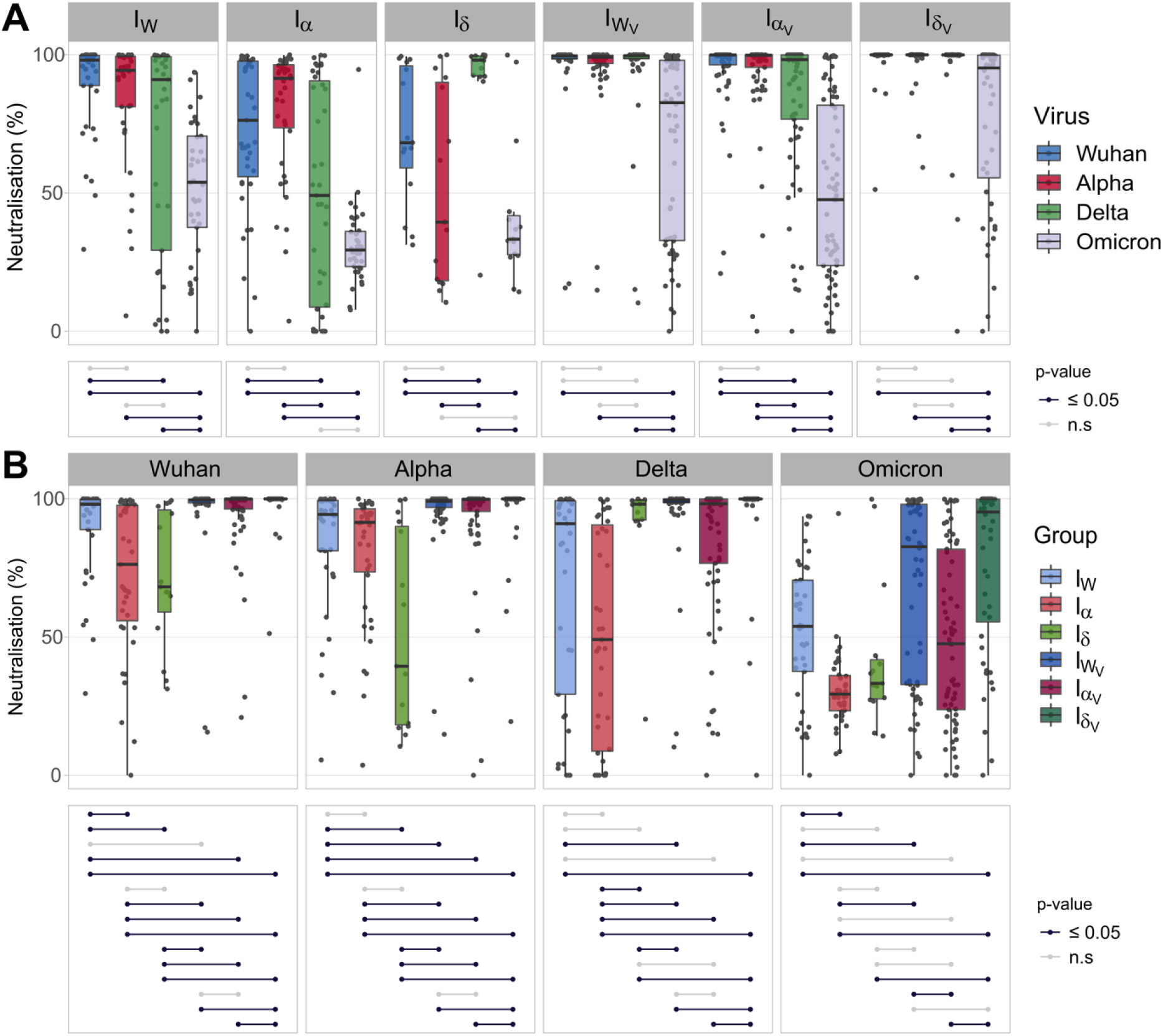
Neutralising responses elicited against pseudotyped viruses carrying the S protein of different SARS-CoV-2 variants according to patient exposure history. (A) Sera from patients were grouped based on immunological histories taking into account the infecting SARS-CoV-2 variant (I_W_: infected with SARS-CoV-2_W_ (light blue), Iα: infected with SARS-CoV-2α (light red), Iδ: infected with SARS-CoV-2δ (light green), I_WV_ infected with SARS-CoV-2_W_ and vaccinated (dark blue), Iα_V_: infected with SARS-CoV-2α (dark red) and vaccinated, Iδ_V_: infected with SARS-CoV-2δ and vaccinated, (dark green)). Neutralising activity was measured using Wuhan (blue), Alpha (red), Delta (green) and Omicron (grey) spike glycoprotein bearing HIV(SARS-CoV-2) pseudotypes and plotted per patient group (A) and per SARS-CoV-2 S variant (B). Neutralisation was measured at a fixed dilution (1:50). Each point represents the mean of two replicates. Boxplots displayed the interquartile range and median values. Significance levels between patient groups or pseudotyped viruses were tested using pairwise Wilcoxon test, and are shown in bottom panels as connected dotplots.

**Figure 6.**
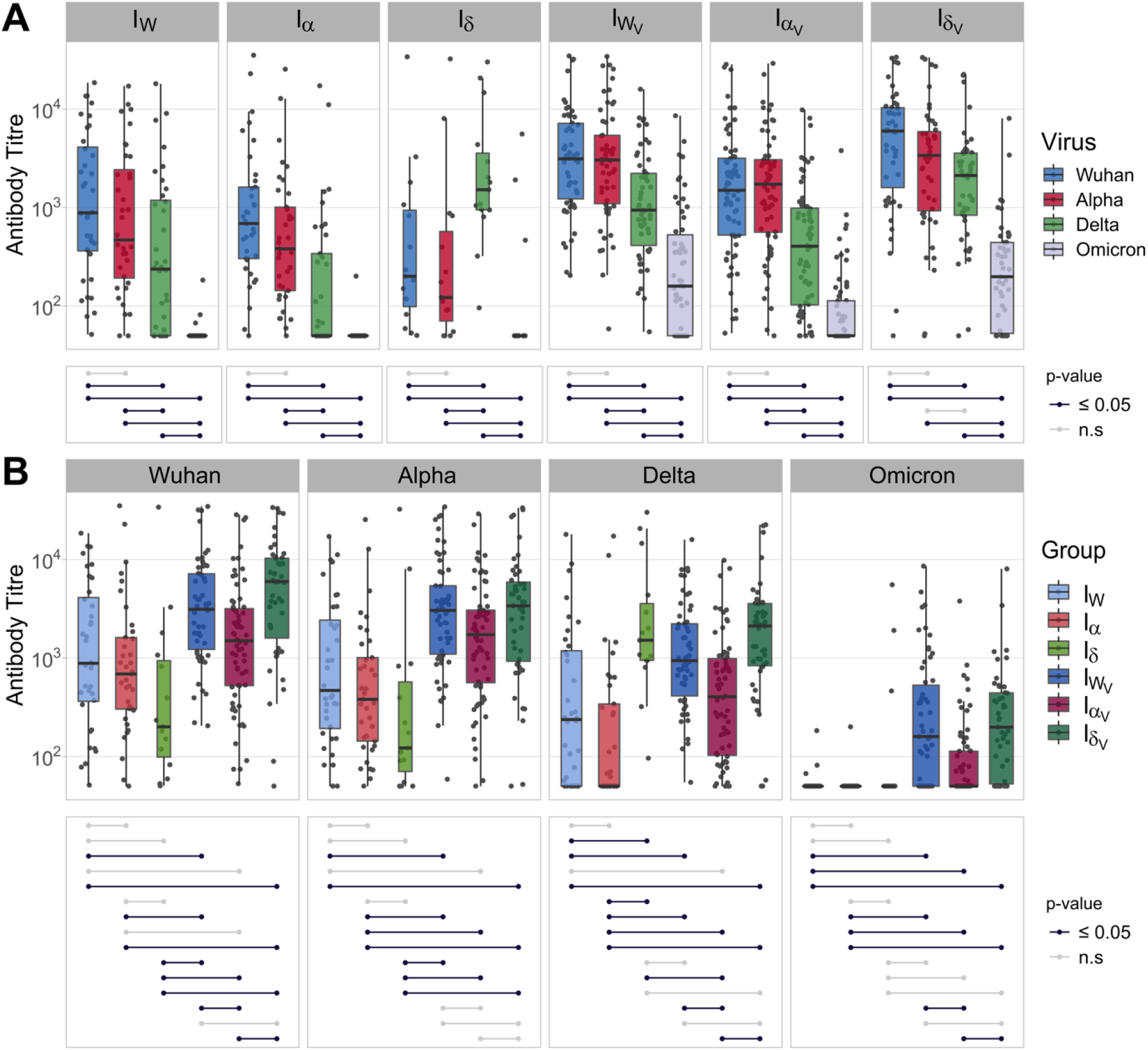
Neutralising antibody titres against SARS-CoV-2 variants in sera from patients with different histories of SARS-CoV-2 exposure taking into account the infecting SARS-CoV-2 variant (I_W_: infected with SARS-CoV-2_W_ (light blue), Iα: infected with SARS-CoV-2α (light red), Iδ: infected with SARS-CoV-2δ (light green), I_WV_ infected with SARS-CoV-2_W_ and vaccinated (dark blue), Iα_V_: infected with SARS-CoV-2α and vaccinated (dark red), Iδ_V_: infected with SARS-CoV-2δ and vaccinated (dark green)). Neutralising activity was measured using Wuhan (blue), Alpha (red), Delta (green) and Omicron (grey) spike glycoprotein bearing HIV(SARS-CoV-2) pseudotypes and plotted per patient group (A) and per SARS-CoV-2 S variant (B). Each point represents the mean of three replicates. Boxplots displayed the interquartile range and median values. Significance levels between patient groups or pseudotyped viruses were tested using pairwise Wilcoxon test, and are shown in bottom panels as connected dotplots.

As the I_V_ group exhibited the highest antibody titres against all variants, we wanted to determine if the order in which patients were exposed to SARS-CoV-2 (either vaccination first or infection first) played any role in the breadth and potency of antibody mediated neutralisation. To test this, we focused on the Iα_V_ group, which exhibited a similar number of patients that had been either infected first (n=28) or vaccinated first (n=41). Sera from patients that had been infected first and then vaccinated exhibited significantly higher neutralisation efficiency against every variant (Fig 7A) and also higher titres of neutralising antibodies (Fig 7B). Overall, this result highlights that the type of exposure (vaccination or infection) and the order in which different types of exposure occur have a significant impact on the breadth and potency of humoral immunity against SARS-CoV-2.

**Figure 7.**
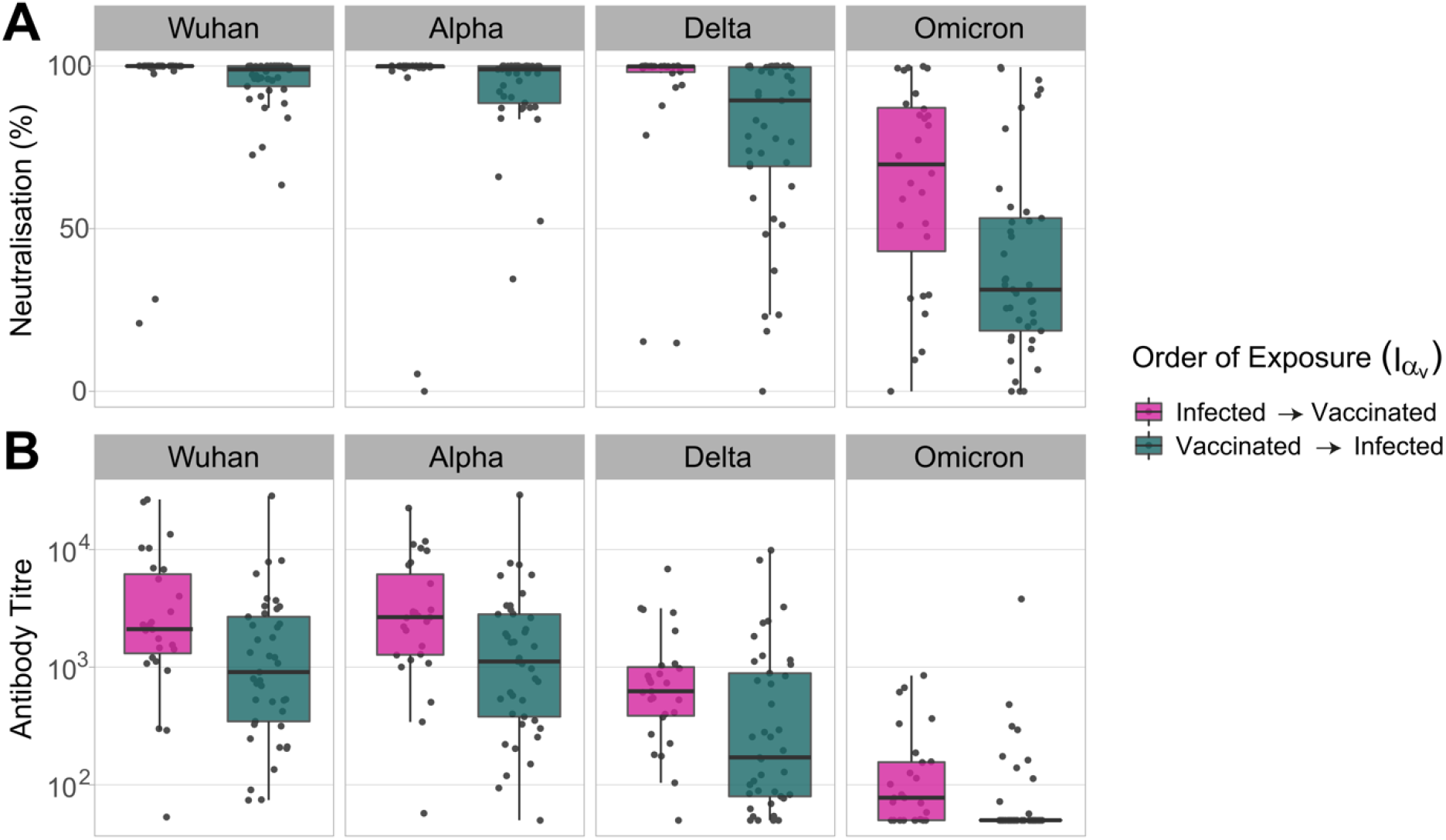
Neutralising responses against pseudotyped viruses carrying the S protein of different SARS-CoV-2 variants were measured (A) and titrated (B) in patients that had been either infected and vaccinated (pink) or vaccinated and infected (seagreen), with SARS-CoV-2α. Serum samples were subject to neutralisation assays using lentiviruses pseudotyped with the S protein of SARS-CoV-2_W_ (Wuhan) SARS-CoV-2_α_ (Alpha), SARS-CoV-2δ (Delta) or SARS-CoV-2ο (Omicron). (A) Neutralisation was measured at a fixed dilution (1:50) and each data point is the mean of two replicates. (B) Antibody titres were calculated by interpolating the point at which infectivity had been reduced to 50% of the value for the non-serum control samples. Each point represents the mean of three replicates.

## Discussion

Our study shows that the immunological landscape of SARS-CoV-2 is highly heterogeneous and has been shaped by the complex interplay between host immunity and virus evolution. Infection by, or vaccination against, SARS-CoV-2 does not elicit lifelong immunity[16, 17] but instead result in a variety of immune phenotypes, which are likely to influence both transmission dynamics and disease outcomes. We demonstrate that multiple factors influence the breadth and potency of the antibody-mediated response against SARS-CoV-2 and include antigenicity of the exposing pathogen, number of exposures, and exposure type (infection and/or vaccination). While T-cell responses play an important role in SARS-CoV-2 immunity[18], we could not evaluate the impact of cellular-mediated immunity due to the nature of our samples (i.e., sera). In line with previous studies, we show that mutations that appeared during SARS-CoV-2 evolution reduce antibody mediated neutralisation[19, 20], suggesting that evolution of the spike gene of SARS-CoV-2 is directional and driven by immune selection. This is consistent with reports of reinfections by novel variants[21, 22]. Our results showing that all serum samples exhibited lowest neutralising activity against pseudoviruses carrying the spike glycoprotein of SARS-CoV-2ο (Figs 4A and 6A) support this view. As all currently licenced vaccine preparations express the S glycoprotein of SARS-CoV-2_W_, it is expected that the risk of reinfections in vaccinated-only individuals increases as SARS-CoV-2 evolves. Similarly, for infected-only individuals, the risk of reinfection will likely increase as the antigenic distance between the viruses involved in primary and secondary infection increases, and thus is a function of time. Our results also show that more exposure events result in broader and more potent antibody-mediated responses (Figs 3B and 5B), and this protective effect is also influenced by the antigenic nature of the viruses involved in the primary infection (or vaccination) and subsequent infections. This finding suggests that updates of the vaccine strains (or development of multivalent vaccines) will improve protection against evolving variants, and also that increased transmission of antigenically divergent SARS-CoV-2 viruses among previously exposed individuals will result in future higher levels of protection. We also show that primary infection followed by vaccination results in more potent humoral responses (Fig 7), which indicates that the type and order of exposure events have a significant impact on the breadth and potency of the antibody mediated response. These findings are consistent with recent reports [13, 14, 23]. While it is not advisable to promote the acquisition of SARS-CoV-2 immunity by natural infection given the risk of severe disease and/or death due to COVID-19 in naive individuals, our results suggest that vaccines based on live-attenuated viruses might provide increased protection.

In sum, our work underscores the complexity of the immunological landscape of SARS-CoV-2. While our results will inform the development of better epidemiological models to predict the future transmission dynamics of SARS-COV-2[24], further clinical studies are needed to determine the impact of exposure history on disease presentation to prepare better for the future disease burden of COVID-19 as this disease becomes endemic.

## Materials and Methods

### Ethics statement

Ethical approval was provided by NHSGGC Biorepository (application 550).

### Serum samples

Random residual biochemistry serum samples (~41,000) from primary (general practices) and secondary (hospitals) healthcare settings were collected by the NHSGGC Biorepository between the 31st of March 2020 and 22nd of September 2021. Associated metadata included date of sample collection, date of positive PCR result, date of first and second vaccination and vaccine manufacturer. Seronegative samples were selected based on their ELISA results (SARS-CoV-2 S1 or SARS-CoV-2 RBD) and the absence of a positive PCR test result or record of vaccination. Samples from vaccinated patients were selected based on their ELISA result (SARS-CoV-2 S1 or SARS-CoV-2 RBD), record of vaccination with 1 or 2 doses at least 14 days prior to blood collection and absence of a positive PCR test result for at least 14 days after sampling. Samples from infected patients had no record of vaccination, were ELISA positive (SARS-CoV-2 S1 or SARS-CoV-2 RBD) and had a positive PCR test result at least 14 days before sample collection. Samples from infected and vaccinated patients were identified by their positive ELISA result, the presence of positive PCR test result and record of vaccination with 1 or 2 doses at least 14 days prior to blood collection. Samples from infected and infected and vaccinated patients were further stratified to infecting variants (based on the date of the positive PCR test), by identifying key time periods during which each variant was most predominant. All serum samples were inactivated at 56°C for 30 minutes before being tested.

### Cells

HEK293T and 293-ACE2 cells were maintained at 37°C, 5% CO_2_, in Dulbecco’s modified Eagle’s medium (DMEM) supplemented with 10% foetal bovine serum, 2mM L-glutamine, 100μg/ml streptomycin and 100 IU/ml penicillin. HEK293 cells were used to produce HEK293-ACE2 target cells by stable transduction with pSCRPSY-hACE2 and were maintained in complete DMEM supplemented with 2μg/ml puromycin. HEK293T cells were used for the generation of HIV(SARS-CoV-2) pseudotypes.

### IgG quantification

IgG antibodies against SARS-CoV-2 the spike and nucleocapsid proteins were measured using an MSD V-PLEX COVID-19 Coronavirus Panel 2 (K15369) kit. Multiplex Meso Scale Discovery electrochemiluminescence (MSD-ECL) assays were performed according to the manufacturer’s instructions. Briefly, 96-well plates were blocked at room temperature for at least 30 minutes. Plates were then washed; samples were diluted 1:5000 and added to the plates along with serially diluted reference standard and serology controls. Plates were incubated for two hours and further washed. SULFO-TAG detection antibody was added, and plates were incubated for one hour. After incubation, plates were washed and read using a MESO Sector S 600 plate reader. Data were generated by Methodological Mind software and analysed using MSD Discovery Workbench (v4.0). Results were normalised to standard(s) and expressed as MSD arbitrary units per ml (AU/ml).

### Neutralisation assays

Pseudotype-based neutralisation assays were carried out as described previously[11]. HEK293T cells were transfected with the appropriate SARS-CoV-2 Spike gene expression vector (Wuhan, Alpha, Delta, or Omicron) together with p8.9171 and pCSFLW72 using polyethylenimine (PEI, Polysciences, Warrington, USA). HIV (SARS-CoV-2) pseudotype-containing supernatants were harvested 48 hours post-transfection, aliquoted and frozen at −80°C prior to use. Gene constructs bearing the Wuhan (D614G), Alpha (B.1.1.7), Delta (B.1.617.2) and Omicron (B.1.1.529) Spike genes were based on the codon-optimised spike sequence of SARS-CoV-2 and generated by GenScript Biotech. Constructs bore the following mutations relative to the Wuhan-Hu-1 sequence (GenBank: MN908947): Wuhan(D614G) – D614G; Omicron (BA.1, B.1.1.529) - A67V, Δ69-70, T95I, G142D/Δ143-145, Δ211/L212I, ins214EPE, G339D, S371L, S373P, S375F, K417N, N440K, G446S, S477N, T478K, E484A, Q493R, G496S, Q498R, N501Y, Y505H, T547K, D614G, H655Y, N679K, P681H, N764K, D796Y, N856K, Q954H, N969K, L981F; Alpha (B.1.1.7)-L18F, Δ69-70, Δ144, N501Y, A570D, P681H, T716I, S982A, D1118H; Delta (B.1.617.2)-T19R, G142D, Δ156-157, R158G, L452R, T478K, D614G, P681R, D950N.

Neutralisation efficiency was measured first using a fixed dilution of serum samples in duplicates. Samples with neutralising activity ≥ 50% relative to the no serum control were then titrated by serial dilutions. Each sample was serially diluted in triplicate from 1:50 to 1:36450 in complete DMEM, incubated for 1 hour with HIV (SARS-CoV-2) pseudotypes, and plated onto HEK239-ACE2 target cells. After 48 hours, luciferase activity was measured by adding Steadylite Plus chemiluminescence substrate and analysed using a Perkin Elmer EnSight multimode plate reader. Antibody titres were estimated by interpolating the point at which infectivity had been reduced to 50% of the value for the no serum control samples. Samples that did not have an antibody titre were arbitrary assigned a value of 50 (the lowest dilution available).

### Statistical analysis

Shapiro-Wilk tests were performed to assess data homoscedasticity. As data were found not to be normally distributed, non-parametric pairwise Wilcoxon Rank Sum tests were carried out to assess statistically significant differences in antibody levels between groups and viruses. Holm’s method was used to adjust p-values to account for multiple statistical comparisons. Separate tests were performed for each group when comparing between viruses, and for each virus when comparing between groups. Pairwise comparisons were presented as connected dotplots, highlighting the significance levels of each of the paired comparisons. All analyses and data visualisations were executed using the stats[25] and ggplot2[26] packages respectively, from R version 4.0.5.

## Data availability

Data of each sample, including metadata and results of each assays, are included in S1 Appendix.

## Acknowledgements

We thank Massimo Palmarini for scientific advice and the NHS Greater Glasgow and Clyde Biorepository for providing serum samples.

## Supporting Information

**S1 Figure.**
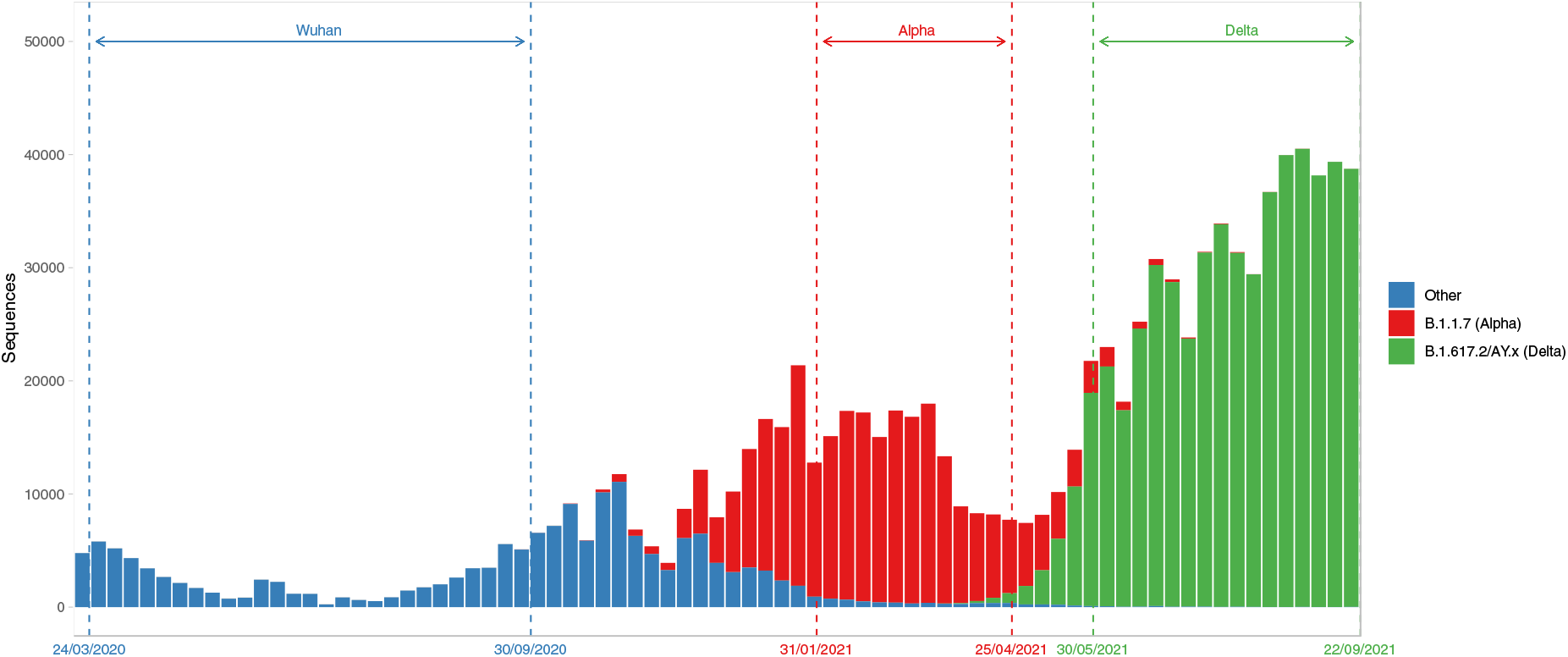
Number of SARS-CoV-2 variants sequenced in the United Kingdom between March 2020 and September 2021. Variants are colour-coded, and the key is shown on the right of the graph. Vertical dashed lines show the time periods used to select samples as described in the main text and methods. **S1 Appendix.** Assay results and metadata of each serum sample used in this study.

